# Accurate genetic profiling of anthropometric traits using a big data approach

**DOI:** 10.1101/033134

**Authors:** Oriol Canela-Xandri, Konrad Rawlik, John A. Woolliams, Albert Tenesa

## Abstract

Genome-wide association studies (GWAS) promised to translate their findings into clinically beneficial improvements of patient management by tailoring disease management to the individual through the prediction of disease risk^1,2^. However, the ability to translate genetic findings from GWAS into predictive tools that are of clinical utility and which may inform clinical practice has, so far, been encouraging but limited^1,2^. Here we propose to use a more powerful statistical approach that enables the prediction of multiple medically relevant phenotypes without the costs associated with developing a genetic test for each of them. As a proof of principle, we used a common panel of 319,038 SNPs to train the prediction models in 114,264 unrelated White-British for height and four obesity related traits (body mass index, basal metabolic rate, body fat percentage, and waist-to-hip ratio). We obtained prediction accuracies that ranged between 46% and 75% of the maximum achievable given their explained heritable component. This represents an improvement of up to 75% over the phenotypic variance explained by the predictors developed through large collaborations^3^, which used more than twice as many training samples. Across-population predictions in White nonBritish individuals were similar to those of White-British whilst those in Asian and Black individuals were informative but less accurate. The genotyping of circa 500,000 UK Biobank^4^ participants will yield predictions ranging between 66% and 83% of the maximum. We anticipate that our models and a common panel of genetic markers, which can be used across multiple traits and diseases, will be the starting point to tailor disease management to the individual. Ultimately, we will be able to capitalise on whole-genome sequence and environmental risk factors to realise the full potential of genomic medicine.

---

Phenotypic prediction of complex traits from genomic data could transform clinical practice by enabling tailored treatment and targeted disease screening programs based on the genetic make-up of the individual, and by facilitating more efficient allocation of resources within the health systems^5–7^. Ultimately, it would help to understand the underlying disease mechanisms and open the targeted search of specific solutions based on this knowledge. With this in mind, large efforts and investments in the past years have been directed towards generating genotypic and phenotypic data for identifying individual genetic variants associated with different traits through genome-wide association studies (GWAS)^8^. Although using this approach a large number of susceptibility variants for many diseases have been identified, the strategy has several limitations. First, the accuracy of prediction has been disappointingly low for traits affected by a large numbers of susceptibility variants^7^. Second, the approach of identifying one single nucleotide polymorphism (SNP) at a time and including such newly identified SNPs in the prediction models as and when they are identified is unpractical if one wishes to use genetic tests for multiple traits because the composition of each trait’s genetic test would need to be continuously updated and each trait would require its own SNP panel. Third, statistical considerations and simulation studies have shown that the accuracy of prediction for complex traits increases by modelling all available SNPs simultaneously^9^.

Recent studies have shown that SNP arrays containing common genetic variants capture a substantial amount of the genetic variation for each trait and that the contributing SNPs have effects generally too small to be detected with current GWAS sample sizes due to the stringent genome-wide significance levels applied^3,10,11^. Furthermore, we have previously shown through simulations that the size of the studies that have estimated heritability from SNP arrays have been too small to properly estimate SNP effects for accurate phenotypic prediction^12^. However, the availability of large genotyped cohorts for which individual-level data is available, e.g. the UK Biobank^4,13^, combined with new and powerful computational tools^12^ capable of fitting complex statistical models to big datasets and access to high-performance computational infrastructure has the potential to provide accurate SNP effects for genomic prediction.

We show that modelling individual-level data of circa 120,000 individuals can lead to accurate predictions across multiple traits by jointly fitting the SNPs of a single array of common SNPs. These predictions significantly improve on the accuracies of models derived from summary statistics obtained from large GWAS meta-analyses and in turn would ease clinical implementation and direct-to-consumer genetic testing, as well as improve the accuracy of the predictions as the sample sizes of the training datasets increase.

We first focused on human height, a highly heritable quantitative trait commonly used as a model in the study of the genetic architecture of complex traits^3,10,14^ and one of the traits for which most contributing loci have been identified to date. To increase the generality of our findings, we then selected four obesity related traits - BMI, percentage body fat, waist-to-hip ratio (WHR) and basal metabolic rate (BMR). In order to jointly estimate the additive effects of all SNPs we fitted them as random effects in a Mixed Linear Model (MLM) on the training population, with gender and age as fixed effects (Online Methods). As the computational requirements of MLM fitting rapidly increases with incrementing sample sizes, we used DISSECT^12^ (https://www.dissect.ed.ac.uk), a software specifically designed to perform genomic analysis in large supercomputers. Each analysis required ~1h of computing time on the ARCHER supercomputer, harnessing the joint power of 1,152 processors. Using the estimated SNP effects to predict the genetic value of individuals in an independent validation dataset (Online Methods), we computed the prediction accuracy as the correlation between these predicted genetic values and the phenotypes corrected for gender and age.

For our analyses, we used the 152,736 genotyped individuals available from the UK Biobank cohort^4^. After applying stringent quality control criteria, we divided our sample into White-British (123,847 individuals) and non White-British (27,685 individuals), the latter including individuals from different ethnic backgrounds (Online Methods and Supplementary Fig. 1). We divided the White-British further into a group of 114,264 unrelated individuals with a relatedness below 0.0625 (i.e. less related to each other than second cousins once removed), another group of 9,583 individuals that had at least one relationship above 0.0625 with the unrelated White-British group, and a group of self-reported White-British (Online Methods and Supplementary Fig. 1). We modelled 319,038 common SNPs, that is, variants with a minor allele frequency (MAF) >0.05 that passed our genotype quality control.

**Figure 1:**
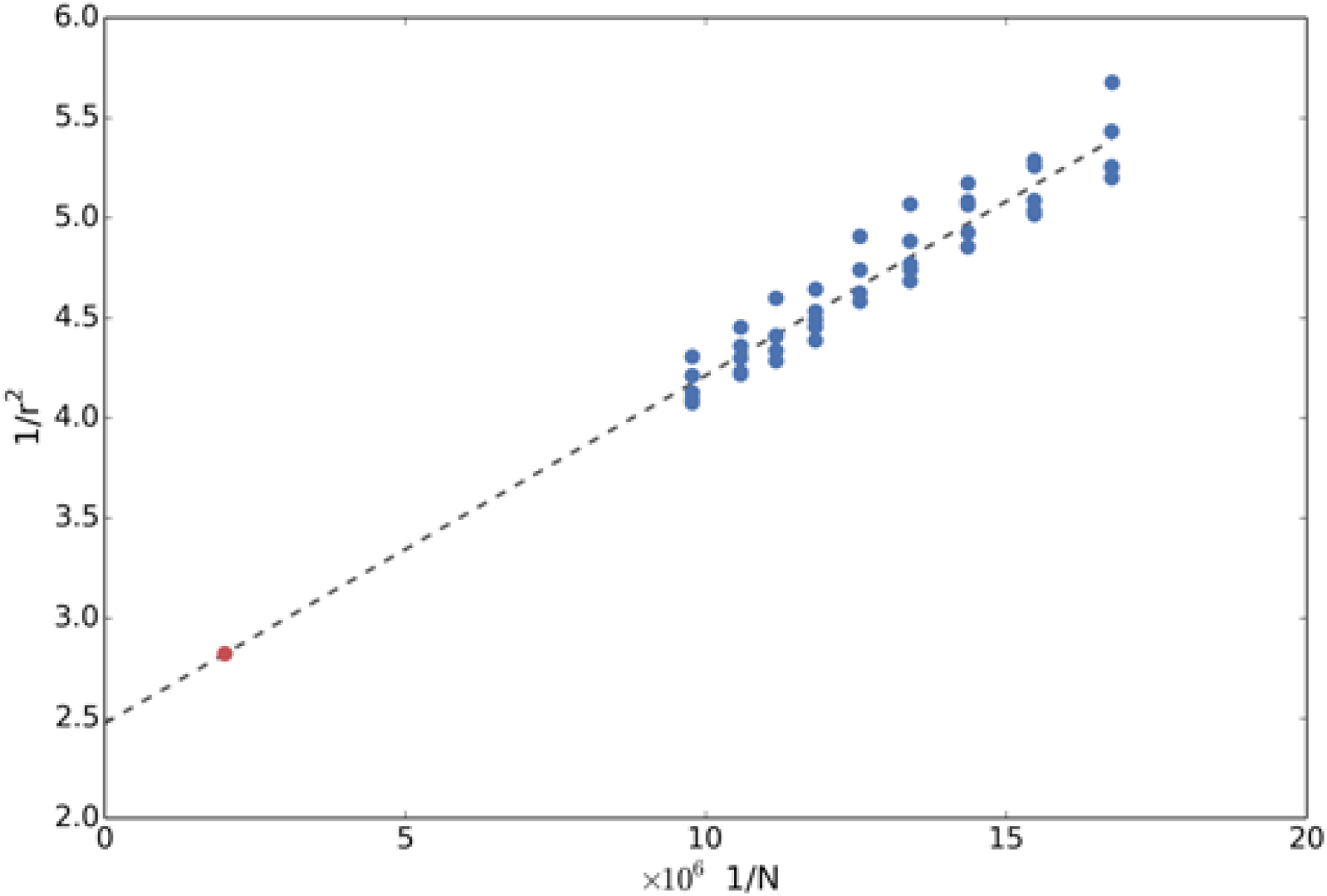
Prediction accuracy as a function of sample size for height. Inverse of the square of the prediction accuracy as a function of the inverse of the training sample size. Blue dots indicate prediction accuracies achieved on several trials. The dashed straight line shows the linear regression fit to the blue dots. The regression intercept indicates the maximum accuracy achievable using common variants represented in the array. The red dot is the expected prediction accuracy with a training sample size of 500,000 individuals.

We used the 114,264 unrelated White-British individuals to train the prediction models and assessed the validity of the within-population predictions using the 9,583 related White-British individuals and the 12,640 self-reported White-British individuals. Prediction accuracy in the self-ported White-British ranged from 0.51 (95% CI 0.49–0.52) for height to 0.20 (95% CI 0.19–0.22) for WHR (Table 1). We evaluated whether prediction accuracies can be further improved by using more complex models (Online Methods). The accuracy for height improved to 0.55 (95% CI 0.53–0.56) despite small reduction in the estimate of heritability. However, accuracies for the other traits decreased. The accuracies we obtained represent between 75% and 46% of the maximum achievable given the estimated SNP-based heritabilities of the traits in unrelated White-British, i.e., 0.53 (SE = 0.004), 0.26 (SE = 0.005), 0.26 (SE = 0.005), 0.20 (SE = 0.005) and 0.31 (SE = 0.005) for height, body fat percentage, BMI, WHR, and BMR, respectively (Online Methods). As expected, phenotypic prediction of relatives was more accurate than that of unrelated people because their phenotypes and genotypes are more correlated to the training samples. The robustness of our within-population predictions were confirmed using 10-fold cross-validation^15^ within the White-British participants (Supplementary Table 1). The SNP effects are available as Supplementary Information.

**Table 1:** Prediction accuracies on related White-British and self-reported White-British.

| Traits | self-reported White-British<br>(95% CI) | related White-British<br>(95% CI) |
| --- | --- | --- |
| Height | 0.51 (0.49-0.52) | 0.53 (0.52-0.55) |
| Body fat percentage | 0.27 (0.25-0.29) | 0.28 (0.26-0.30) |
| BMI | 0.25 (0.24-0.27) | 0.27 (0.26-0.29) |
| WHR | 0.20 (0.19-0.22) | 0.23 (0.21-0.25) |
| BMR | 0.32 (0.31-0.34) | 0.34 (0.32-0.36) |

We also investigated to what extend across-population predictions were feasible. To this end, we further subdivided the non White-British subset by self-reported ethnic background. Excluding ethnicities with less than 1,000 individuals and removing outliers resulted in 7,541 White individuals who did not self-report as White-British, 1,954 Asian or Asian-British individuals, and 1,591 Black or Black-British individuals (Online Methods). Predictions obtained in the White cohort (Table 2) were almost as accurate as to those obtained in the self-reported White-British cohort. Predictions for the other two ethnicities remained considerable but lower than within-population predictions, especially for Black or Black British as expected from the genetic distance between populations (Supplementary Fig. 2), indicating that predictions may benefit from within-ethnic group tailored models.

**Table 2:**
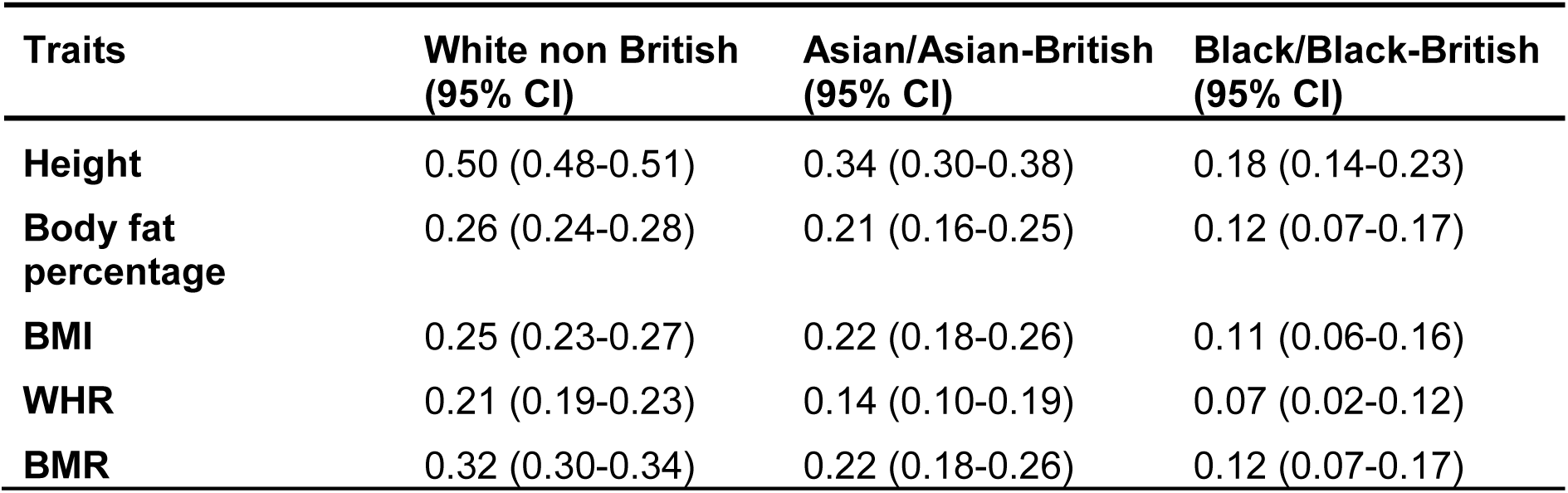
Across-population prediction accuracies.

| Traits | White non British<br>(95% CI) | Asian/Asian-British<br>(95% CI) | Black/Black-British<br>(95% CI) |
| --- | --- | --- | --- |
| Height | 0.50 (0.48-0.51) | 0.34 (0.30-0.38) | 0.18 (0.14-0.23) |
| Body fat percentage | 0.26 (0.24-0.28) | 0.21 (0.16-0.25) | 0.12 (0.07-0.17) |
| BMI | 0.25 (0.23-0.27) | 0.22 (0.18-0.26) | 0.11 (0.06-0.16) |
| WHR | 0.21 (0.19-0.23) | 0.14 (0.10-0.19) | 0.07 (0.02-0.12) |
| BMR | 0.32 (0.30-0.34) | 0.22 (0.18-0.26) | 0.12 (0.07-0.17) |

Although samples sizes for training the models will increase in the future, it is unlikely that they will increase indefinitely. Therefore, we argued that it would be useful to know what sample size would be required to exploit all the genetic variation captured by the SNP array. To gauge that, we computed prediction accuracies for samples of decreasing size, by randomly subsampling the unrelated White-British individuals (Online Methods). Our data fitted very well to a well-known theoretical model^16^. Our predictions suggest that prediction accuracies for height will reach 0.6 (SE = 0.02) when training the models using ~500,000 individuals (Fig. 1), the samples planned to be genotyped UK Biobank in the near future. This prediction accuracy would represent 82% of the maximum accuracy possible given the explained heritability. Similarly, we estimate that genetic prediction models for BMI, WHR, body fat percentage and BMR will reach prediction accuracies of 0.36 (SE = 0.05), 0.29 (SE = 0.03), 0.37 (SE = 0.03), and 0.42 (SE = 0.03) respectively (Supplementary Fig. 3).

Our results confirm previous findings that many variants with small effect can explain a large proportion of the genetic variance. Due to several factors, this part of the genetic variance has so far remained largely unexploited for phenotypic prediction. These factors include the statistical methods used, the available sample sizes, and computational software available to analyse the data. However, as we have shown, predictions which are significantly more accurate can be obtained by increasing sample sizes and using powerful computational approaches to jointly estimate all SNP effects. The phenotypic variance explained by our predictor for height is ~75% larger than that of the largest height meta-analysis to date, which used a discovery sample size ~250% larger than ours, but that used genome-wide significant SNPs from the meta-analyses as predictors^3^. Our prediction accuracies are very close to the maximum achievable given the estimates of the corresponding explained heritable components, and we predict that they will become even closer when the number of samples increase (e.g. when the UK Biobank is fully genotyped). For BMI, which is affected by a smaller genetic component, our predictor explains slightly more variance than previous work that used a discovery sample size ~3 times larger^17^. Finally, we demonstrated that more complex models have the potential to further improve prediction accuracies, although our results also indicate that the optimal model may be trait specific. In conclusion, the presented results support our initial hypothesis and suggest a promising future for genomic prediction of complex traits.

## Methods

### Genotype Quality Control

For our analysis, we used the data for the genotyped individuals in phase 1 of the UK Biobank genotyping program. 49,979 individuals were genotyped by using the Affymetrix UK BiLEVE Axiom array and 102,750 individuals by using the Affymetrix UK Biobank Axiom array. Details regarding genotyping procedure and genotype calling protocols are provided elsewhere (http://biobank.ctsu.ox.ac.uk/crystal/refer.cgi?id=155580). From the overlapping markers, we excluded those which were multi-allelic, their overall missingness rate exceeded 2% or they exhibited a strong platform specific missingness bias (Fisher’s exact test, P < 10^−100^). We also excluded individuals if they exhibited excess heterozygosity, as identified by UK Biobank internal QC procedures (http://biobank.ctsu.ox.ac.uk/crystal/refer.cgi?id=155580), if their missingness rate exceeded 5% or if their self-reported sex did not match genetic sex estimated from X chromosome inbreeding coefficients. These criteria resulted in a reduced dataset of 151,532 individuals. Finally, we only kept the common variants (i.e. with a MAF > 0.05) and those that did not exhibit departure from Hardy-Weinberg equilibrium (P < 10^−50^) in the unrelated (subset of individuals with a relatedness below 0.0625) White-British cohort (see below).

### Ethnicity

The UK Biobank samples are from individuals of diverse ethnicities. To define the White-British cohort, we performed a Principal Components Analysis (PCA) of all individuals passing genotypic QC using a linkage disequilibrium (LD) pruned set of 99,101 autosomal markers (http://biobank.ctsu.ox.ac.uk/crystal/refer.cgi?id=149744) that passed our SNP QC protocol. The related and unrelated White-British individuals were defined as those for whom the projections onto the leading twenty genomic principal components fell within three standard deviations of the mean and who identified themselves as White-British. We defined the other removed White-British as self-reported White-British. The other ethnicities were defined using the self-identified ethnic background (http://biobank.ctsu.ox.ac.uk/crystal/field.cgi?id=21000). As we did with White-British individuals, we only retained those individuals whose projections onto the leading twenty genomic principal components fell within three standard deviations of the ethnicity group mean (Supplementary Fig. 2).

### Phenotype quality control

We defined outliers as males and females that were outside ±3 standard deviations from their gender mean of all the individuals in the UK Biobank, and removed them from the analyses.

### Software

The genotype quality control and data filtering was performed using plink^18^ (https://www.cog-genomics.org/plink2). The PCA, MLMs fittings for estimating SNP effects and phenotype predictions were performed using DISSECT (https://www.dissect.ed.ac.uk) on the UK National Supercomputer (ARCHER). DISSECT software is designed to perform genomic analyses on very large sample sizes without the need to perform mathematical approximations by using the power of large supercomputers.

### Phenotype prediction

The effect of all SNPs were estimated together as a random effect using the model,

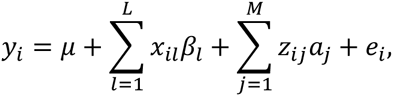
where *μ* is the mean term and *e_i_* the residual for individual *i. L* is the number of fixed effects, *x_il_* being the value for the fixed effect *l* at individual *i* and *β_i_* the estimated effect of the fixed effect *l. M* is the number of markers and *z_ij_* is the standardised genotype of individual *i* at marker *j*. The vector of random SNP effects a is distributed as 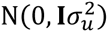. The vector of environmental effects e is distributed as 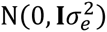. The heritabilities were estimated as 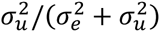.

The prediction of the phenotype 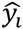 for the individual *i* was computed as a sum of the product of the SNP effects and the number of reference alleles of the corresponding SNPs:

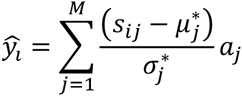

Where *s_ij_*· is the number of copies of the reference allele at SNP *j* of individual *i, M* is the number of SNPs used for the prediction, and *a_j_* the effect of SNP *j*. 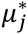 and 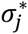 are the mean and the standard deviation of the reference allele in the training population.

Prediction accuracies were computed as the correlation between the predicted phenotype and the real one after correcting by the estimated effect of the used covariates (e.g. sex and age).

### Phenotype prediction using a two variance components model

The MLM of the previous section assumes that all SNP effects follow a Gaussian distribution with one variance. However this is may not be true. To improve the model we first fitted all SNPs independently using a standard GWAS model,

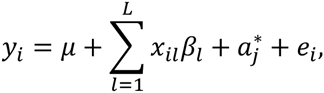

Here, the parameters are the same as in the previous MLM, and the SNP effect size 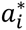 is estimated independently for each SNP as a fixed effect. We then divided the SNPs into two groups based on their effect size. Specifically, one group of SNPs in the main distribution and a group of outliers, which were defined as SNPs with effect sizes more than 3 standard deviations away from the mean effect across all SNPs. Using these groups, we fit an extended MLM where we assume the SNP effects were distributed in two different Gaussian distributions with a different variance each one,

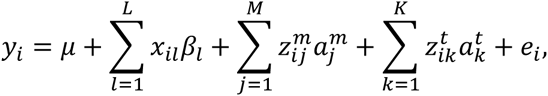
where all parameters are the same as in the simpler MLM, but now M and K are the number of SNPs in the main distribution and the two tails, respectively, while 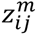 and 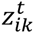 are the corresponding genotypes. We fit independent variances for the two groups of SNPs, so that the vector of SNP effects in the main distribution, a*^m^*, is distributed as 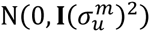 and the vector of SNP effects in the tails, a*^t^*, is distributed as 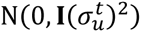.

### Random subsampling

We computed accuracies for samples of decreasing size, by randomly subsampling 5 of the 10-fold cross-validation subsets used in the within unrelated White-British population predictions (Supplementary Table 1).

## Supplementary Information

Supplementary Information contains the SNP estimated effect sizes for each trait.

## Acknowledgments

This work was mainly supported by the Medical Research Council [grant numbers MR/K014781/1 and MR/N003179/1] and The Roslin Institute Strategic Grant funding from the BBSRC. AT also acknowledges funding from the Medical Research Council Human Genetics Unit. This work used the ARCHER UK National Supercomputing Service (http://www.archer.ac.uk) and the Edinburgh Compute and Data Facility (ECDF) (http://www.ecdf.ed.ac.uk/). This research has been conducted using the UK Biobank Resource. We also acknowledge the fruitful comments received from Chris Haley, Wendy Bickmore and Pau Navarro.

## Contributions

All authors contributed equally on this study.

## Ethical approval

The use of the UK Biobank dataset falls within the study’s ethical approval from the North West Medical Research Ethics Committee (Reference 11/NW/0382). An informed consent was obtained from all subjects.

## URLs

DISSECT and documentation available at: https://www.dissect.ed.ac.uk

PLINK2 and documentation available at: https://www.cog-genomics.org/plink2

